# Long-term effect of forest harvesting on boreal species assemblages under climate change

**DOI:** 10.1101/2022.08.21.504664

**Authors:** Ilhem Bouderbala, Guillemette Labadie, Jean-Michel Béland, Junior A. Tremblay, Yan Boulanger, Christian Hébert, Patrick Desrosiers, Antoine Allard, Daniel Fortin

## Abstract

Logging is the main human disturbance impacting biodiversity in forest ecosystems. However, the impact of forest harvesting on biodiversity is modulated by abiotic conditions through complex relationships that remain poorly documented. Therefore, the interplay between forest management and climate change can no longer be ignored. Our aim was to study the expected long-term variations in the assemblage of bird and beetle communities following modifications in forest management under different climate change scenarios. We developed species distribution models to predict the occurrence of 87 species of birds and beetles in eastern Canadian boreal forests over the next century. We simulated three climate scenarios (baseline, RCP4.5 and RCP8.5) under which we varied the level of harvesting. We also analyzed the regional assemblage dissimilarity by decomposing it into balanced variations in species occupancy and occupancy gradient. We predict that forest harvesting will alter the diversity by increasing assemblage dissimilarity under all the studied climate scenarios, mainly due to species turnover. Species turnover intensity was greater for ground-dwelling beetles, probably because they have lower dispersal capacity than flying beetles or birds. A good dispersal capacity allows species to travel more easily between ecosystems across the landscape when they search for suitable habitats after a disturbance. Regionally, an overall increase in the probability of occupancy is projected for bird species, whereas a decrease is predicted for beetles, a variation that could reflect differences in ecological traits between taxa. Our results further predict a decrease in the number of species that increase their occupancy after harvest under the most severe climatic scenario for both taxa. We anticipate that under severe climate change, increasing forest disturbance will be detrimental to beetles associated with old forests but also with young forests after disturbances.

## 1 Introduction

Biodiversity is facing an unprecedented rate of change due to human activities (Sala et al., 2000; Newbold et al., 2015). For terrestrial ecosystems, land-use changes through forest harvesting, agriculture or urbanization are considered the main causes of biodiversity losses and reductions in ecosystem services (Sala et al., 2000; Naeem et al., 2012; Bichet et al., 2016; Matuoka et al., 2020). Human exploitation of natural resources can induce animal and plant mortality, force species displacement, increase biotic interactions, modify their life-history traits, change their morphology and physiology, and even their polymorphic forms (Sergio et al., 2018). Moreover, the exploitation of natural habitats by humans results in habitat losses and biotope fragmentation, which constrains the size of animal and plant populations (Coelho et al., 2020). For instance, by returning stands to an early successional stage, forest harvesting homogenizes forest landscape, which results in an overrepresentation of early-succession forests at the expense of the older states (Cyr et al., 2009; Boucher et al., 2017). Forest harvesting targets old-stands, which are economically more profitable but these stands are not homogeneous as they present structural diversity and can be distinguished into different groups Martin et al. (2018). Moreover, these stands are the habitat of a biodiversity indicator species, the Black-backed Woodpecker *Picoides arcticus*), associated with old-growth attributes. Thus, forest harvesting may threaten vertebrate species associated with old-growth forest attributes Martin et al. (2021) but also organisms (insects, fungi, non-vascular plants) that have low-dispersal capacity and for show extinction risk increases rapidly when ecological continuity is broken Nordén and Appelqvist (2001).

Climate change is the second most important driver of change in biodiversity after land use, for the terrestrial ecosystems (Sala et al., 2000; Dobson et al., 2021). Climate change may impact forest landscapes indirectly by triggering natural disturbances such as wildfire, which strongly alter forest age structure and composition (Boulanger et al., 2014, 2017; Boulanger and Puigdevall, 2021), and to which are associated a unique biodiversity. Climate change may also directly impact tree growth and alter competitive abilities of boreal tree species Boulanger and Puigdevall (2021). These alterations are predicted to be detrimental to biodiversity especially those associated with older mixedwood and coniferous forests (Tremblay et al., 2018; Cadieux et al., 2020). It may also impact species directly by changing their occurrence or abundance, when climatic conditions go out of their tolerance limits, due to modifications in temperature and precipitation conditions (Micheletti et al., 2021; Bouderbala et al., 2022), which might reshape community composition by species turnover (Bouderbala et al., 2022). For example, the displacement of species to higher elevations is one of the predicted consequences of climate change to overcome the potential increase in temperature (Davis and Shaw, 2001). However, climate change could have implications on the abundance of species with low dispersal capacity that are unable to follow the climate conditions to which they are currently adapted (Davis and Shaw, 2001; Muluneh, 2021) and could lead to an increase in extinction risks (Thomas et al., 2004).

Although modelling biodiversity changes is becoming essential to predict the effects of human activities on the global environment, many challenges remain because there are many related issues that need to be integrated (Ceaus, u et al., 2021). There are urgent needs to identify the most appropriate forest harvesting scenarios to achieve management objectives, such as species recovery Cadieux et al. (2019) or habitat restoration (Jones et al., 2022). This would help identify more targeted conservation actions under climate change such as vegetation restoration.

Despite the number of studies made to better understand the cumulative effects of the different types of disturbances on biodiversity (Sala et al., 2000), there are considerable uncertainties over the magnitude of changes in future animal communities Almeida-Rocha et al. (2020). The cumulative effect of anthropogenic disturbances could induce complex effects on biodiversity through the adjustment in species’ ecological traits, which may have cascading repercussions from individual changes to community-level responses (Sergio et al., 2018). However, most literature evoking the modelling of biodiversity changes considered the assumption of no interactions among the various drivers (Sala et al., 2000). In this work, we investigate how the interaction between forest harvesting and climate change would impact biodiversity. Specifically, we examine future implications of forest harvesting levels on the variation of bird and beetle assemblages in Québec’s boreal forest under different climate change scenarios. We aim first to quantify the extent of assemblages’ alteration by increasing harvest levels under different climate change scenarios and second to determine the main drivers. Moreover, we study how species associated with different types of habitat classes will be affected by the combined effects of forest harvesting and wildfire.

## 2 Materials and methods

### 2.1 Study area and occurrence data

The study area was located in the Côte-Nord region of Québec, Canada (48°*N* to 53°*N*, 65°*W* to 71°*W*) within an area of 114,118 km^2^ (Fig. 1). This region is part of the eastern spruce-moss subdomain and is dominated by black spruce (*Picea mariana* [Mill.] BSP) with balsam fir (*Abies balsamea* [L.] Miller). Forest harvesting had been the main source of anthropogenic disturbances since the late 1990s (Bouchard and Pothier, 2011). Moreover, This area is mainly affected by two major natural disturbances: spruce budworm outbreak (SBW) (*Choristoneura fumiferana*) and frequent wildfires (Boucher et al., 2017; Labadie et al., 2021).

**Figure 1:**
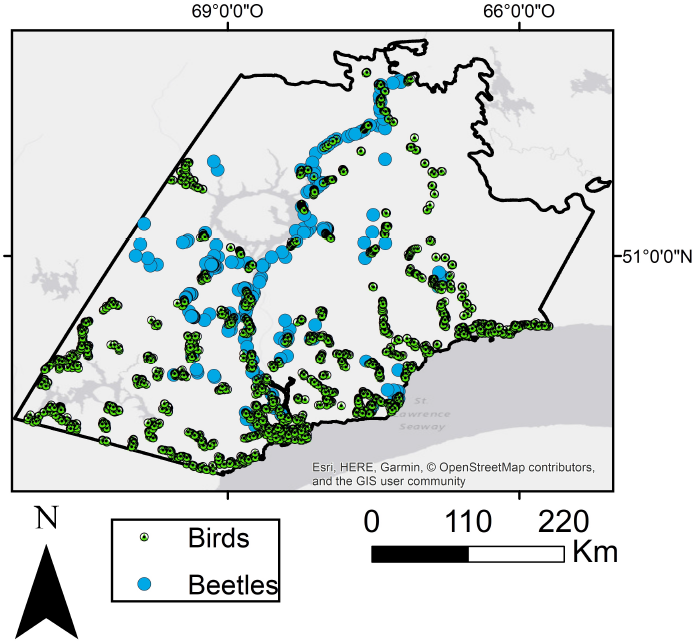
Study area location of forest stands sampled for birds and beetles.

We used presence-absence data collected between 2004 and 2018 to model the species distribution of two taxa: birds and beetles. For birds, we used the data from the Second Atlas of Breeding Birds of Québec (Atlas des oiseaux nicheurs du Québec, 2018), based on species occurrence detected using unlimited distance 5-minute point counts (Bibby et al., 2000) collected during the breeding season, from late May to mid-July. For beetles, we merged different databases collected in 2004, 2005, 2007, and 2011 (Janssen et al., 2009; Légaré et al., 2011; Bichet et al., 2016). We added 54 sites sampled in 2018 in the northern part of the territory, along the northeast principal road going to Labrador. Insect sampling involved one multidirectional flight-interception trap per site to capture the flying beetles, and four meshed pitfall traps per site to sample beetles that move on the soil surface (Janssen et al., 2009; Bichet et al., 2016). Data from the sites heavily impacted by the SBW outbreak (Ministère des Forêts de la Faune et des Parcs, 2018) were removed to focus only on the effect of the wildfires. We used for this purpose a cumulative index of defoliation severity (Labadie et al., 2021) (we removed 4% and 7% of sampling sites for beetle and bird databases, respectively).

### 2.2 The forest management scenarios

Three harvest levels were considered: no harvest activity (NoHarvest), 8% of the areas harvested over 10 years (Harvest) – the typical harvest rate for the study area – and half this level (i.e., 4%, Harvest_0.5_). The harvest levels were combined with three climate scenarios: baseline level, radiative forcing associated with the Representative Concentration Pathway scenario (RCP) 4.5, which involved a 3°C increase in mean temperature from 2000 to 2100; and RCP 8.5 associated with a 7.5°C increase over the same period. We compared the response of species occurrence and occupancy between different harvest intensities under three climate scenarios, which explains why we had three reference scenarios (Table 1). Also, we took into consideration in our simulations the northern limit of territorial forest attributes of harvest activities. Beyond this limit, no forest management was authorized by the Ministry of Forests, Wildlife, and Parks (Ministère des Forêts de la Faune et des Parcs, 2018).

**Table 1:** The studied scenarios. We considered three of them for reference (Climate=Baseline, Harvest=NoHarvest; Climate=RCP 4.5, Harvest=NoHarvest; Climate=RCP 8.5, Harvest=NoHarvest) and compared them to six other scenarios. Under each climate scenario (Baseline, RCP 4.5 and RCP 8.5) we compared NoHarvest to Harvest_0.5_ and Harvest, which makes six comparisons in total.

| Reference scenario |  | Simulated scenario |  | Compared landscapes |
| --- | --- | --- | --- | --- |
| Climate | Harvest | Climate | Harvest | (Reference-Simulated) |
| Baseline | NoHarvest | Baseline | Harvest<sub>0.5</sub> | Baseline (NoHarvest-Harvest<sub>0.5</sub>) |
| Baseline | NoHarvest | Baseline | Harvest | Baseline (NoHarvest-Harvest) |
| RCP 4.5 | NoHarvest | RCP 4.5 | Harvest<sub>0.5</sub> | RCP 4.5 (NoHarvest-Harvest<sub>0.5</sub>) |
| RCP 4.5 | NoHarvest | RCP 4.5 | Harvest | RCP 4.5 (NoHarvest-Harvest) |
| RCP 8.5 | NoHarvest | RCP 8.5 | Harvest<sub>0.5</sub> | RCP 8.5 (NoHarvest- Harvest<sub>0.5</sub>) |
| RCP 8.5 | NoHarvest | RCP 8.5 | Harvest | RCP 8.5 (NoHarvest-Harvest) |

### 2.3 Forest landscapes simulation

We simulated forest succession across landscapes using LANDIS-II (Scheller and Mladenoff, 2004), a forest landscape model that captures forest succession across landscapes as an emergent property of both stand and landscape-scale processes (Scheller et al., 2007). Specifically, we used the LANDIS-II Biomass Succession extension v5.0, Base Fire v4.0, BDA v4.0, together with the Biomass Harvest v5.0 (Scheller and Mladenoff, 2004) to simulate forest succession, fire (Scheller and Domingo, 2005) and spruce budworm outbreak disturbances as well as harvesting in each 250-m cell at a 10-year time step. Outbreaks of SBW were simulated using the LANDIS-II Biological Disturbance Agent (BDA) extension v3.0 (Sturtevant et al., 2004, 2012). The Biomass Succession extension simulates modifications in cohort aboveground biomass (AGB) over time by taking into consideration cohort age, life-history traits, and land type responses of individual tree species. A random forest model was used to predict crown closure covariate by using Canadian National Forest Inventory (NFI) forest attribute maps (Beaudoin et al., 2014). These maps are a k-Nearest Neighbours interpolation of the NFI photoplot data acquired in 2001 and are depicting over 130 forest attributes including species-specific biomass, stand age and crown closure at a 250-m resolution (see Beaudoin et al., 2014). We, therefore, build a random forest model predicting cell-level crown closure in NFI products from NFI species-specific biomass as well as stand age. This model had a very high goodness of fit (*R*^2^ = 0.86). The model was then applied on LANDIS-II outputs to predict crown closure all along the simulation, for each cell by using simulated species-specific biomass and stand age (Labadie et al., 2022). Using species group and predicted crown closure, we created five land cover classes from the Earth Observation for Sustainable Development of Forests (EOSD) Land Cover Classification Legend (Beaubien et al., 1999; see Table 2). See (Labadie et al., 2022) for more details regarding the spatially-explicit forest simulation model with LANDIS-II.

**Table 2:** Natural habitats used (if less than 50% of the pixel’s surface is in disturbance).

| Habitat | Condition |
| --- | --- |
| Dense conifer | - vegetation exceeds more than 50% of the surface<br>- canopy density is higher than 60% of the surface<br>- coniferous stands occupy more than 75% of the surface |
| Open conifer | - vegetation exceeds more than 50% of the surface<br>- canopy density is less than 60% of the surface<br>- coniferous stands occupy more than 75% of the surface |
| Mixed-wood | - vegetation exceeds more than 50% of the surface<br>- neither coniferous nor broad-leaf trees account for more than 75% of the surface<br>- the vegetation surface with trees is greater than the surface without trees |
| Open habitat | - vegetation exceeds more than 50% of the surface<br>- the vegetation surface with trees is less than the surface without trees |
| Other | - Non-vegetation (trees, shrubs, grasses, bryophytes) that exceeds more than 50% of the surface |

### 2.4 The predictor variables

Land cover associated with sampling sites was determined from the Canadian National Forest Inventory referenced in the year 2001 (Beaudoin et al., 2014), updated yearly for subsequent fires and cuts. The landscape was split into 11 land cover types. Specifically, we considered land covers with stand age greater than 50 years, i.e., closed-canopy conifer forest, open-canopy mature conifer forest, mixed forest, and open area (see Table 2), together with six disturbance classes, i.e, fire and forest harvesting subdivided by age classes: 0–10, 11–20 and 21–50 years (Labadie et al., 2021). The category of the other types of habitat was not included in the models to avoid collinearity. In the same region of study and forest types, Janssen et al. (2009) showed that ground-dwelling and flying beetles were influenced by landscape variables within radius around sample plots of 400 and 800 m respectively, while Bichet et al. (2016) used a 1 km radius for birds. Therefore, we used for ground-dwelling beetles 20 pixels (231 m resolution) centered on the focal pixel (i.e., 21 pixels in total) to calculate the frequencies of the land cover classes with a radius of 2.5 pixels (*∼*0.58 km). For flying beetles and birds, we used 37 pixels with a radius *∼*0.81 km to calculate land cover frequencies. We also considered the distance to the nearest burned areas as well as the stand age as potential predictor variables.

### 2.5 Association between bird species and habitat type

We considered three land cover classes to calculate species winning conditional probability (WCP), i.e., an increase in the probability of species occupancy from uncut to post-harvest in 2100, conditional to species habitat (see Eq. 2). We calculated WCP for (1) generalist bird species; (2) species associated with early-to-mid succession forest (EMSF), which included the following land covers: wetland, deciduous, mixed-wood (young to mid-age), and coniferous habitat (young to mid-age); and (3) species associated with late succession forest (LSF), which included the following land covers: mixed-wood (mature to old), deciduous (mature to old) and coniferous habitat (mature to old). Stand age was classified as follows: young (stand age < 30 years), mid-age (stand aged between 30 and 60 years), mature (stand aged between 60 and 80 years), and old (stand age greater than 80 years). This classification was used only for birds because information regarding beetles’ associated land cover was unavailable at the moment of the study. Bird-associated land cover has been based on habitat associations from Gauthier and Aubry (1995) and Robert (2019).

### 2.6 Relationship between beetles and forest disturbances

A literature review was done on every beetle taxon to characterize two important functional traits: habitat preference and dispersal capacity. First, we identified species that respond positively to forest harvesting or burned forests, the remainder being considered as species associated with undisturbed natural forests. Then, in order to discriminate beetle taxa that mainly disperse by flight from those that mainly walk on the forest floor, we calculated abundance ratios from the data collected. For each beetle taxon, we pooled the total abundance (from 2004 to 2018) per flight interception trap and divided by the total abundance per pitfall trap. This ratio provides an index of beetle proficiency to fly or to walk on the ground. As the ratio provides continuous estimates, we kept only beetles with a ratio higher than 2 as flying beetles and beetles with a ratio lower than 3 as ground-dwelling beetles. Beetle species with ratios between 2 and 3 were removed from the study.

### 2.7 Modelling framework

We developed species distribution models (SDMs) (Guisan and Zimmermann, 2000; Guisan and Thuiller, 2005; Peterson et al., 2011) to estimate single-species occurrence probability based on extensive field surveys of 88 species (39 flying beetles, 15 ground-dwelling beetles and 34 bird species). The methodology here is described in details inBouderbala et al., 2022 as Habitat-Only-Based Models (HOBMs). We used generalized linear mixed models to predict species occurrence probability (package ‘lme4’; Bates et al., 2015). A random intercept was added to include any differences between sampling years based on three full potential models deferring only in the fixed effects of the models. We partitioned the modelling strategy into several steps starting from data pre-processing to the species occurrence probability projections. SDMs were estimated only for species recorded at least at 1% and 5% of sampling sites for birds and beetles, respectively. We used land cover variables extracted from LANDIS forest landscape simulations for the simulated scenarios (see Table S1 for predictor variables description). We also standardized the predictor variables just before any model calibration to facilitate the convergence of the models (MacKenzie et al., 2017). We removed highly correlated variables (|*r*| > 0.60, where *r* represented pairwise Pearson correlation coefficient) and kept the 5 most important potential predictor variables according to the Akaike information criterion (AIC) calculated on a univariate generalized linear model (GLM) with linear and quadratic terms for each variable (Zurell et al., 2020; Bouderbala et al., 2022).

We suggested three full potential models, the first one contained only linear forms of the predictor variables. However, the second and third full models included also quadratic transformations of the covariates (see Table S2). The best models were selected for each full model based on a 10-fold cross-validation procedure by minimizing the AIC criterion (package ‘MuMIn’; Barton, 2015). For each species, we kept only the models with *AUC* ≥ 0.7 (Araújo et al., 2005; Hosmer et al., 2013). In addition, we analyzed the assemblage structure based on continuous occurrence probabilities instead of binarizing the outputs (Guisan and Thuiller, 2005; Grenié et al., 2020).

We calculated the percentage change in the regional occupancy probability (ROP) between the reference (*Ref*) and the simulated (*Sim*) scenarios 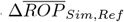 (Bichet et al., 2016) as follows:

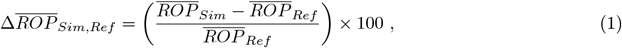

where 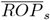 denotes the average of the ROP over all the species under the scenario *s* and is given by

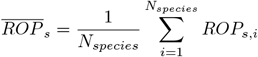

With

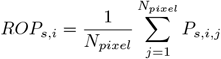

and where *N*_*species*_, *N*_*pixel*_ and *P*_*s,i,j*_ respectively represent the number of species, the number of pixels in the study area, and the probability of occurrence of the species *i* for the scenario *s* at the cell *j*.

Winning conditional to habitat or to relation to disturbance probability was calculated as

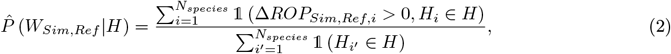

where 1 is the indicator function and *H*_*i*_ is the associated habitat for bird species *i* (*H* ∈ {G, EMSF, LSF}) or the relation to disturbance for beetle species (*H* ∈ {Fire, Harvest, Fire − Harvest, None}). In addition, we used the regional Bray-Curtis dissimilarity index (*RBC*_*Sim,Ref*_) to measure the degree of dissimilarity in assemblage composition between the compared scenarios (Baselga, 2010, 2013). The *RBC*_*Sim,Ref*_ between the scenarios *Sim* and *Ref* was defined as

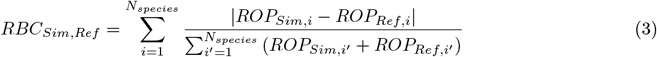

We split the Bray-Curtis dissimilarity index into two additive components:

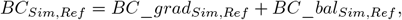

where *BC*_*grad*_*Sim,Ref*_ is the occupancy gradient and *BC*_*bal*_*Sim,Ref*_ is the species balanced variation in occupancy. The latter was defined as

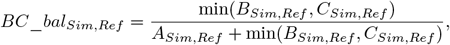

Where

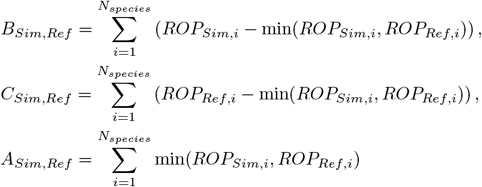

To quantify the main source of assemblage dissimilarity, we performed a decomposition of the regional Bray-Curtis dissimilarity, based on the probability of occurrence, into *balanced variation in species occupancy* and *occupancy gradient*, which are the generalizations of species turnover and the nestedness respectively in the context of continuous output (Baselga, 2013) and occurrence probabilities in our case (Bouderbala et al., 2022). This decomposition of the dissimilarity measure computed based on scenarios helped us to characterize the mechanisms that drive variation in species composition. Hereafter the terms species turnover and nestedness will be used as equivalents of *balanced variation in species occupancy* and *occupancy gradient* respectively. We then computed the ratio

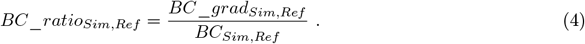

The case *BC*_*ratio*_*Sim,Ref*_ > 0.5 then indicated that the community change was mostly caused by nestedness, whereas *BC*_*ratio*_*Sim,Ref*_ < 0.5 indicated the dominance of species turnover in the assemblage dissimilarity. We used the package ‘betapart’ (Baselga and Orme, 2012) for the assemblage analysis.

## 3 Results

Of 61, 105 and 42 of candidate species of birds, flying beetles and ground-dwelling beetles, 34, 39 and 15 were selected for the projection with *AUC* ≥ 0.7 (Fig. 2a-b-c) respectively. Stand age was the most frequently selected predictor variable for the two taxa (Fig. 2d). Moreover, ground-dwelling beetles were more closely associated with old undisturbed forests than flying beetles. However, the response of flying beetles was much greater to the distance from burned forests than ground-dwelling-beetles, which showed a weaker response that decreased with time after wildfire (Fig. 2d).

**Figure 2:**
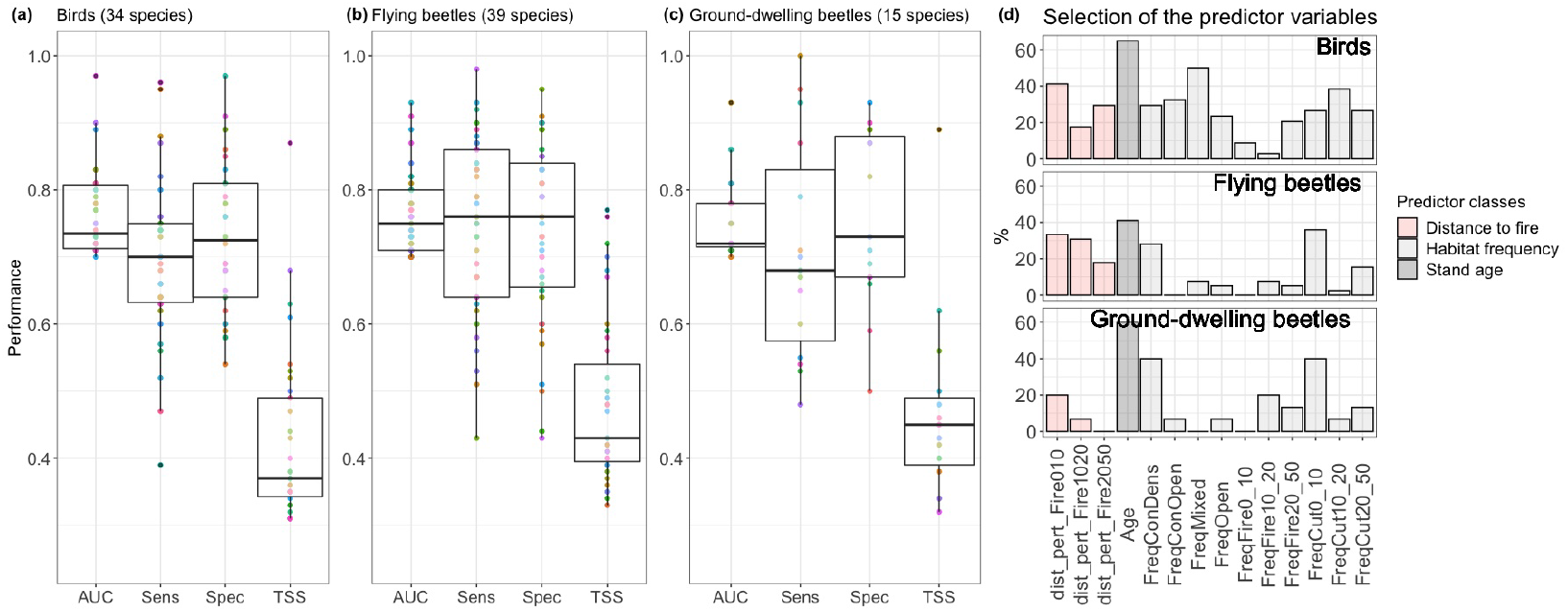
Performance metrics (area under the curve (AUC), sensitivity (Sens), specificity (Spec) and the true skill statistic (TSS)) of the selected models (**a, b, c**). Percentage of the predictor variables that were included in the species regressions (**d**). Abbreviations: distance to the nearest fire (dist_pert_Fire), stand age (Age), frequency of conifer dense (FreqConDens), frequency of conifer open (FreqConOpen), frequency of mixed wood (FreqMixed), frequency of open habitat (FreqOpen), frequency of disturbance by fire (FreqFire), frequency of disturbance by cut (FreqCut).

### 3.1 Changes in disturbance level following climate change

Changing climate from the baseline to RCP 8.5 scenarios in 2100 increased the proportion of burned stands at each harvest scenario. For example, the proportion of burned stands was 17.2% under the baseline-Harvest scenario, 24.7% under RCP 4.5-Harvest scenario, and 31.6% under RCP 8.5-Harvest scenario (Table. 3). Furthermore, the increase either in wildfire or in harvest levels decreased the proportion of coniferous forests (dense, open and mixed wood). Moreover, climate change as well as forest harvesting decreased significantly stand age. For instance, under RCP 4.5 the mean stand age decreased from 90 to 46 years from NoHarvest to Harvest scenarios. Also, the mean stand age under Harvest_0.5_ went from 67 to 49 years from baseline to RCP 8.5 (Table 3).

**Table 3:** Percentage of the area occupied by the different land covers in 2100 (stand age average in years). Abbreviations: conifer dense (ConDens), conifer open (ConOpen), mixed wood (Mixed), open habitat (Open), Other, disturbance by fire (Fire), disturbance by cut (Cut) and Fire+Cut (Disturbance).

| Landscape | ConDens | ConOpen | Mixed | Open | Other | Fire | Cut | Total | Disturbance |
| --- | --- | --- | --- | --- | --- | --- | --- | --- | --- |
| Baseline-NoHarvest | 10.83(156.91) | 18.93(136.15) | 30.97(142.48) | 0.96(0) | 23.18(31.66) | 15.13(24.17) | 0(-) | 100(97.89) | 15.13 |
| Baseline-Harvest<sub>0.5</sub> | 6.91(149.04) | 11.5(123.48) | 22.7(120.13) | 0.96(0) | 23.87(28.84) | 17.5(24.88) | 16.56(22.36) | 100(66.7) | 34.06 |
| Baseline-Harvest | 4.83(146.46) | 8.1(116.74) | 17.83(109.52) | 0.96(0) | 23.87(27.87) | 17.17(20.58) | 27.25(23.25) | 100(52.57) | 44.42 |
| RCP 4.5-NoHarvest | 10.74(161.74) | 17.87(142.92) | 28.59(140.72) | 0.96(0) | 18.9(14.35) | 22.94(17.35) | 0(-) | 100(89.83) | 22.94 |
| RCP 4.5-Harvest<sub>0.5</sub> | 6.66(154.74) | 10.92(130.48) | 21.43(117.39) | 0.96(0) | 20.06(15.83) | 23.76(17.28) | 16.21(23.64) | 100(60.82) | 39.97 |
| RCP 4.5-Harvest | 4.17(152.53) | 6.74(120.14) | 17.01(105.03) | 0.96(0) | 19.85(15.32) | 24.65(19.05) | 26.6(22.49) | 100(46.06) | 51.25 |
| RCP 8.5-NoHarvest | 6.67(160.38) | 11.58(144.26) | 24.68(144.53) | 0.96(0) | 20.36(21.31) | 35.76(15.17) | 0(-) | 100(72.83) | 35.76 |
| RCP 8.5-Harvest<sub>0.5</sub> | 4.24(152.07) | 7.12(133.56) | 17.24(122.12) | 0.96(0) | 20.48(19.13) | 36.32(14.77) | 13.65(21.82) | 100(49.27) | 49.97 |
| RCP 8.5-Harvest | 2.94(151.72) | 5.3(126.17) | 14.65(113.17) | 0.96(0) | 21.14(19.21) | 31.63(15.31) | 23.38(21.48) | 100(41.66) | 55.01 |

### 3.2 Assemblage analysis

Species assemblage dissimilarity was higher in NoHarvest-Harvest than in NoHarvest-Harvest_0.5_, under the three climate scenarios in 2100 (Fig. 3). Under RCP 4.5, Bray-Curtis dissimilarity increased from 0.08 to 0.12 for birds and 0.06 to 0.09 for flying beetles and from 0.08 to 0.11 for ground-dwelling beetles when comparing NoHarvest-Harvest_0.5_ to NoHarvest-Harvest scenarios. Compared to birds and ground-dwelling beetles, flying beetles had lower dissimilarity in assemblages regardless of climate scenarios (Fig. 3). However, the dissimilarity in species assemblage over harvest gradient decreased with climate change (see the difference in Bray-Curtis dissimilarity between baseline and RCP 8.5 in Fig. 3). This result was also observed in the structure of land cover under RCP8.5 compared to RCP4.5 and baseline by varying harvest intensity (see Fig S4).

**Figure 3:**
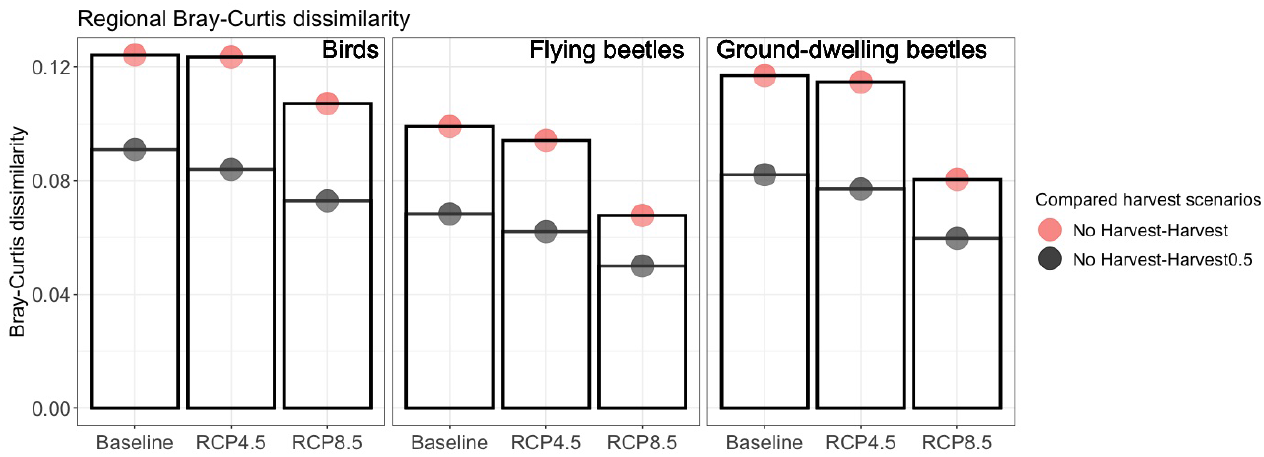
Regional Bray-Curtis dissimilarity measures (Eq. 3) computed based on the difference in species regional occupancy between reference and simulated scenario. At each climate change scenario (Baseline, RCP 4.5, RCP 8.5), we computed the dissimilarity between Harvest_0.5_ and Harvest vs. NoHarvest in 2100. Under Baseline, we compared respectively Baseline-NoHarvest to Baseline-Harvest_0.5_ and Baseline-NoHarvest to Baseline-Harvest. Under RCP 4.5 we compared respectively RCP 4.5-NoHarvest to RCP 4.5-Harvest_0.5_ and RCP 4.5-NoHarvest to RCP 4.5-Harvest. Finally, under RCP 8.5 we compared respectively RCP 8.5-NoHarvest to RCP 8.5-Harvest_0.5_ and RCP 8.5-NoHarvest to RCP 8.5-Harvest. The abbreviation “None” represents the case of species associated to closed forests.

### 3.3 Occupancy analysis

We observed a change in the regional occupancy between the scenarios of forest harvesting (Harvest_0.5_ and Harvest vs. NoHarvest) under all three climate change scenarios. More precisely, we observed an increase in regional occupancy in the bird community from nonharvest to harvest (passage from 13% to 15% in the regional occupancy from nonharvest to harvest scenarios under RCP 4.5). The opposite behaviour was observed in the beetle community (for instance a decrease in regional occupancy from 31% to 27% was observed for flying beetles by comparing the same scenarios). These changes in the overall occupancy of birds, and the decrease for beetles from NoHarvest to Harvest cases was visible over 114,118 km^2^ of the study area (Fig. 4) and also regionally (Fig. 5a). Post-harvesting changes in species assemblages were mainly caused by species turnover (*BC*_*ratio*_*Sim,Ref*_ < 0.5) for the two taxa with a strongest turnover for ground-dwelling than flying beetles. Under each climate scenario, the occupancy of bird species associated with young habitats together with generalist species increased significantly contrary to species associated with mature and old forests, when we compared harvested to unharvested scenarios (Fig. 5b). Under baseline and RCP 4.5, 50% of bird species associated with late succession forest increased in occupancy from no harvested to harvested scenarios (Fig. 6a), whereas 82% of species associated with early to mid succession forest increased in occupancy (see Fig S2 for the variation at the species scale). By comparing RCP 8.5 to RCP 4.5, the proportion of winner species (from non-harvested to harvested landscapes) that were associated with early to mid-succession forests decreased from 82% to 73%. Under the same conditions, species associated with late succession forests increased from 50% to 63% (Fig. 6a). When we compared non-harvested to harvested scenarios for flying beetles, species that were associated with burned stands had the weakest increase in occupancy compared to species associated with harvested stands or closed forests (Fig. 5c).

**Figure 4:**
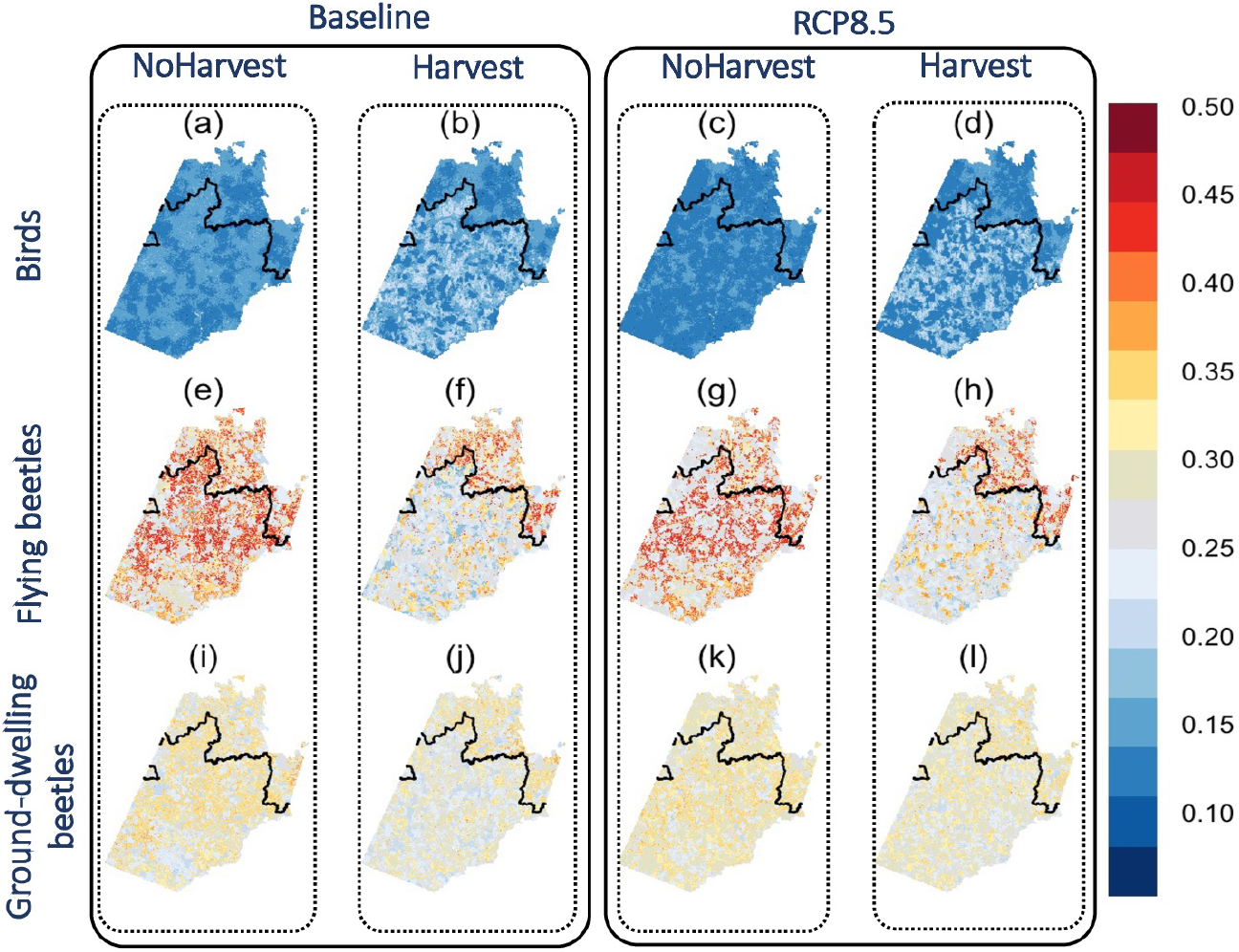
The average maps of the potential occurrence distribution for each taxon based on the four scenarios **BaselineNoHarvest, BaselineHarvest, RCP 8.5NoHarvest** and **RCP 8.5Harvest**. The black line represents the northern limit for forest harvesting activities.

**Figure 5:**
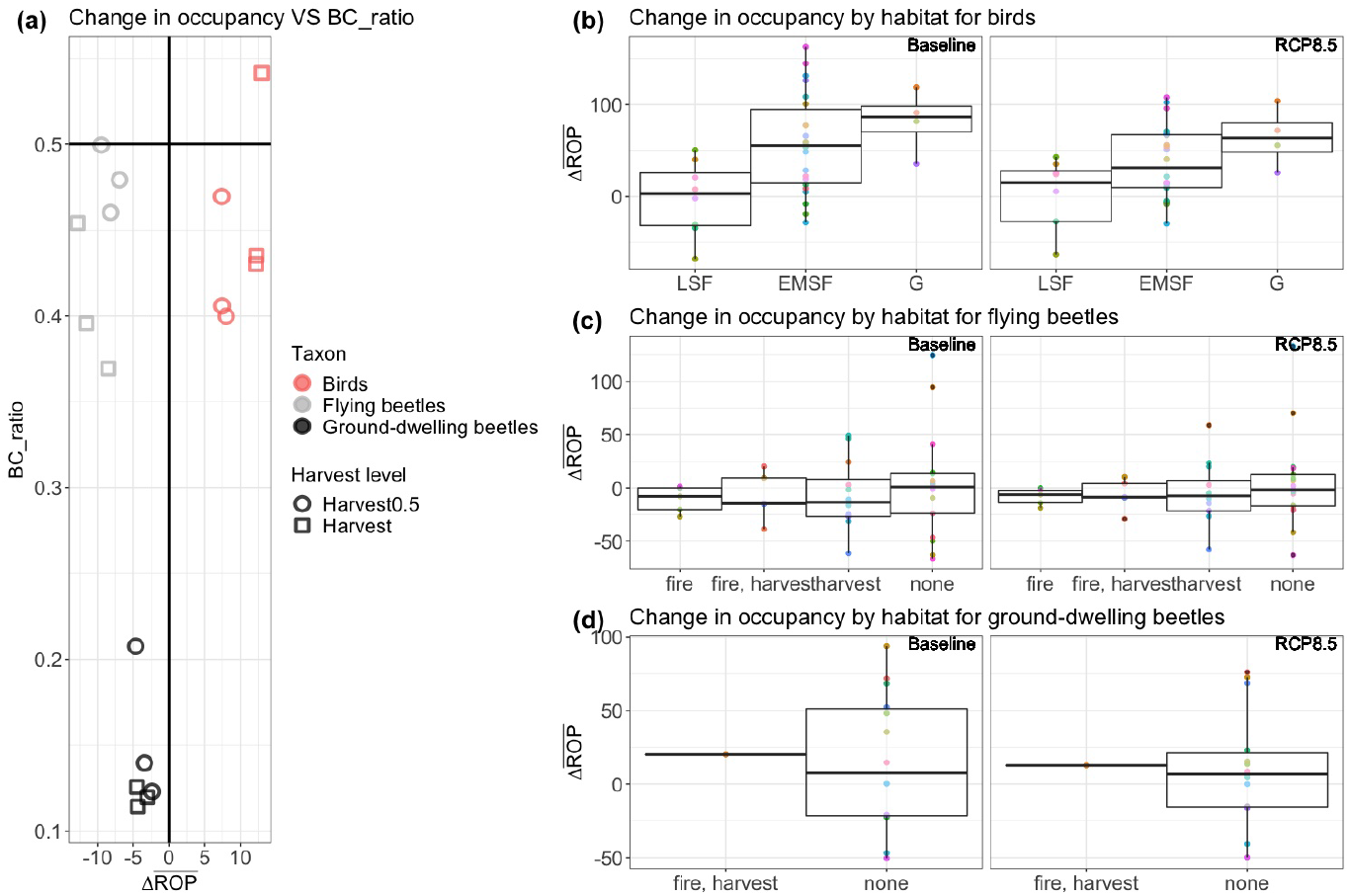
**(a)** Regional Bray-Curtis dissimilarity ratio (*BC*_*ratio*) with the percentage of change in the regional occupancy probability 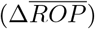. At each climate change scenario (Baseline, RCP 4.5, RCP 8.5), we computed the change in the ROP (Eq. 1) and Bray-Curtis ratio (Eq. 4) between Harvest_0.5_ and Harvest vs. NoHarvest in 2100. **(b)** Percentage of change in the regional occupancy probability between Harvest and Noharvest under the baseline and RCP 8.5 for bird species. **Abbreviations**: late succession forest (*LSF*), early-to-mid succession forest (*EMSF*), generalist species (*G*).**(c-d)** Percentage of change in the regional occupancy probability between Harvest and Noharvest under the baseline and RCP 8.5 for flying and ground-dwelling beetles species with relation to a disturbance.

**Figure 6:**
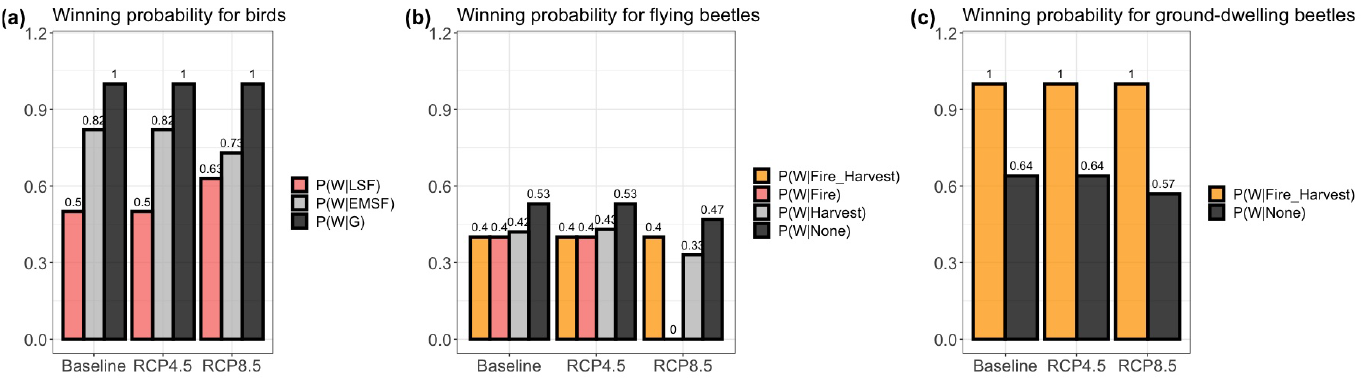
Conditional winning probability (Eq. 2) of bird species among land covers when the landscape changes from NoHarvest to “Harvest”, under the three climates scenarios. Under Baseline, we compared Baseline-NoHarvest to Baseline-Harvest. Under RCP 4.5 we compared RCP 4.5-NoHarvest to RCP 4.5-Harvest. Finally, under RCP 8.5 we compared RCP 8.5-NoHarvest to RCP 8.5-Harvest.

The percentage of winning flying beetles associated with closed forests decreased from 53% to 47% from baseline to RCP 8.5 under a forest harvest scenario (Fig. 6b). Winning species of flying beetles associated with harvested stands even decreased from 42% to 33% under the same conditions. Among flying beetles associated with burned forests, the percentage of winning species went from 40% to 0%. Regarding ground-dwelling beetles, the percentage of winning species associated with closed forests decreased from 64% to 57% under the harvest scenario under climate change (baseline to RCP 8.5) (Fig. 6c).

## 4 Discussion

Forest harvesting can impact significantly forest biota and may have lasting effects on biodiversity (Linden-mayer and Franklin, 2002). We aimed to quantify how the interaction between forest harvesting and climate change would impact biodiversity. We analyzed how species assemblages can be expected to change under different forest management scenarios and climate conditions. Our models suggest that forest harvesting will cause pronounced changes in species assemblages for birds and beetles by 2100 under all climate change scenarios. Our result is supported by many empirical studies (Baker et al., 2016; Bichet et al., 2016) evoking the noticeable effect of harvesting on the assemblage of common species and predicted that the effect could be stronger by including rare species (Bichet et al., 2016). Moreover, Tremblay et al. (2018) projects a drastic reduction in the habitat of the Black-backed Woodpecker (*Picoides arcticus*), a focal species associated with deadwood and representative of old-growth forest biodiversity in eastern Canada (Tremblay et al., 2020). The simulated change in assemblage dissimilarity was mainly dominated by species turnover. This implies that the change in the assemblage composition from NoHarvest to Harvest scenarios was dominated by an increase in occupancy for a subset of species that was balanced mostly by a decrease in occupancy for the other subset of species.

It is important to underline that the effects of forest harvesting were studied under each climate scenario. So, the difference between the less disturbed scenario (baseline-NoHarvest, i.e. 15% of disturbance) and the most disturbed scenario (RCP8.5-Harvest, i.e. 55% of disturbance) on species occurrence could be different from our results based on comparing different harvest levels within the same climate scenario. Our results predict an increase in occupancy in the majority of bird species associated with Early-to-Mid succession forests contrary to species associated with Late succession forests. This result is coherent with previous studies (Fortin et al., 2011; Bichet et al., 2016). However, under the extreme climate scenario, there is one more late-succession species increasing in occupancy when we compared harvested to unharvested scenarios, while this is the opposite for early-to-mid succession species which decreased by one species. This difference could be explained by the proportion of burned stands on the harvestable stands under each climate scenario. Indeed, we observed a decrease in the harvested stands with climate change contrary to the burned stands. Those interactions between forest harvesting and future climate change could impact the occupancy of species that depend on disturbed stands. For example, on the one hand, we observed that Yellow Warbler (YEWA, *Setophaga petechia*), a bird species associated with early-to-mid succession forest and negatively affected by both harvested forest and stand age regressed under RCP 8.5 contrary to the baseline climate scenario. On the other hand, Swainson’s Thrush (SWTH, *Catharus ustulatus*) that was associated with late succession forest and negatively affected by burned forests increased under RCP 8.5 (see Fig. S1). The increase in wildfires under the extreme climate scenario could affect negatively species that prefer harvested stands if forest managers increase salvage logging. It has been shown that bird trait assemblages differ between burned, post-fire salvage, and traditionally logged habitats in North American forests (Bognounou et al., 2021).

Our results on beetles confirm that the dispersal capacity of taxa is an important functional trait to understand future extinction risks for this order of insects. As they are less mobile, ground-dwelling beetles are mainly associated with local conditions (Janssen et al., 2009). In our study, 93% of them were associated with mature and undisturbed forests. Nevertheless, the frequency of young harvested stands had some support in the models, suggesting that ground-dwelling beetle communities found in old boreal forest persist some years in recently harvested stands, as shown for carabids by Niemela et al. (1993) in western Canada. For instance, two common carabids, *Platynus decentis* and *Pterostichus adstrictus*, selected in our models were “winners” under each climate change scenario. Post-disturbance successions are usually characterized by a deciduous stage, which should favor *P. decentis* as it is usually found in deciduous and mixed forest (Work et al., 2008). Moreover, *P. adstrictus* is favored by wildfire (Cobb et al., 2007) and is not affected by forest management (Work et al., 2008). Two species of *Cryptophagus* (Cryptophagidae), a mycetophagous family usually associated with mature and undisturbed forests, also showed a positive response to harvest intensification. They have low dispersal capacities and their persistence in harvested landscapes suggest that suitable conditions could be maintained as the area of study is covered by a matrix of old-growth forests growing in a humid climate that favor fungal growth. Certain ground-dwelling beetles also respond to recently burned stands. For instance, some carabids are known to be attracted to recently burned stands (Holliday, 1991). In a recent study, Bell et al. (2022) suggested that pyrophily in insects have evolved to take advantage of unique post-fire environmental conditions. Indeed, heat-sterilized ovipositing substrates are used by *carabids* of the genus *Sericoda* to increase their reproductive output. These conditions are ephemeral and *Sericoda* disappears rapidly after fire. This could result from suboptimal reproductive conditions. Many other insects are known to respond in such a way to wildfire (Boucher et al., 2016).

We showed that species turnover was the dominant source of dissimilarity in species assemblages for the two studied taxa but ground-dwelling beetles registered the highest rate. This might be due to their lower moving capacity. They cannot escape easily new conditions generated by disturbances. As seen earlier, they can persist for some years after harvesting, but the new habitat conditions end up inducing a higher species turnover. In the same boreal region, Le Borgne et al. (2018) showed that post-logging beetle assemblages during forest succession were mostly driven by interspecific interactions than by habitat attributes. It was probably a consequence of the presence of a pristine forest matrix in the surrounding landscape in which local communities differ as they have evolved together over a long time. Our results on ground dwelling beetle occupancy showed a reduction of the rove beetles (Staphylinidae), which are mainly associated with mature and undisturbed forests. They represent 83% of the ground-dwelling “losers” regardless of the climatic change scenario. As the rove beetles modelled were all predators and thus at the top of the food chain, interspecific interactions are critical, and they might be the first to disappear in such turnovers.

In comparison, beetle communities that fly to colonize burned trees after wildfire are much more structured according to habitat characteristics (Azeria et al., 2012). Among modelled flying beetles in our study, only 44% of species were associated with undisturbed forests. Most were saproxylic beetles associated with burned forests. The time window for successful colonization of burned trees could be narrow for some saproxylic beetle as phloeophagous species may simply benefit from the high-quality sub-cortical tissues of freshly burned trees, usually for 1-2 years only (Muona and Rutanen, 1994; Wikars, 1994; Saint-Germain et al., 2004b,c; Boulanger and Sirois, 2014). After tree felling, woody debris decay for several decades, allowing great persistence for late-succession beetles (Jonsell and Weslien, 2003; Boulanger and Sirois, 2014). The complexity of the different saproxylic beetle feeding guilds is expressed in the analysis of the change in occupancy for flying beetles. Even though 26% were associated with fire, the “winners” among them dropped from 40% under base scenario to 20% under the RCP8.5 scenario. For instance, in the literature, the six bark beetle species modelled were associated with fire, harvest or both disturbances, but were all “losers” under both climate change scenarios. Saint-Germain et al. (2004a) showed the preference of these beetles to colonize large diameter burned trees, showing the importance of this forest attribute for maintaining biodiversity. However, it is probably the first attribute to be negatively affected by harvesting and climate change. Moreover, wildfire (Azeria et al., 2012; Boucher et al., 2016), salvage logging (Norvez et al., 2013) and traditional logging (Légaré et al., 2011) rapidly homogenize stands and biodiversity at the landscape level. On the long-term, this lowers landscape heterogeneity as well as biodiversity at the landscape level. As observed for ground-dwelling beetles, changes in occupancy are also most visible for predators in flying beetles. While the proportion of “losers” increases from 50 to 63% between climate change scenarios for predatory ground dwelling beetles, it increases from 44 to 77% for predatory flying beetles. The scenarios tested integrate burned forests interacting with increased harvesting and the resulting landscapes suggest that an erosion in the flying beetle food chain is likely to impact all trophic levels.

Our results were based only on the most common species, the response of rare specialists and threatened species might decrease the regional occupancy with both harvest activities and climate conditions (Cadieux et al., 2019). Therefore, it will be interesting to analyze the response of the rare species to disturbance. One possible approach could be through the development of a probabilistic model which combines the occurrence probability of rare species with indicator species.

In conclusion, forest harvesting activities are predicted to modify the composition of beetle and bird species assemblages under all climate scenarios. We also indicated that the consequences of maintaining the current level of forest harvesting under the worst climate change scenario could modify the species-habitat association. Ideally, we aim to generate a situation where both early and late successional stages can coexist by managing forests. Interestingly, we observed opposite responses in occupancy between birds, which was generally positive, and beetle, which was negative, in our study area. The perfect separation between the two taxonomic groups regarding the change in occupancy could reflect the impact of ecological traits such as movement capacity and diet differences between boreal species. In any case, forest management must be adapted to achieve conservation objectives under all climate scenarios. Our simulations show that harvest activities must be reduced. Even when it maximizes biodiversity, harvest could be harmful to taxa with low moving capacity (beetles in our case) which characterize old-growth forests.

## Supporting information

**File S1. The studied beetle and bird species**.

**Table S1 Description of the used predictors**.

**Table S2 The potential 3 full models**.

**Fig S1 Change in ROP for bird species with the associated habitat**.

**Fig S2 Change in ROP for flying beetles species with the relation to a disturbance**.

**Fig S3 Change in ROP for ground-dwelling beetles species with the relation to a disturbance**.

**Fig S4 The distribution of the land cover under the simulated management scenarios in 2100**.

## Acknowledgements

This work was supported by the Sentinel North program of Laval University, funded by the Canada First Research Excellence Fund. A.A., D.F., C.H., and P.D. were also supported by The Natural Sciences and Engineering Research Council of Canada (NSERC). We acknowledge Calcul Québec and Compute Canada for their technical support and computing infrastructures. We also thank the Québec Breeding Bird Atlas for supplying data. We would also like to thank the following partners: Regroupement QuébecOiseaux, Environment and Climate Change Canada, and Birds Canada, as well as all of the volunteer participants who gathered data for the project. We are grateful to Louis-Paul Rivest for his valuable statistical suggestions. Special thanks are due to Alexandre Terrigeol for his valuable efforts regarding data collection, as well as to Georges Pelletier and Nicolas Bédard for beetle identification.

