## Supplementary material for "Long-term effect of forest harvesting on boreal species assemblages under climate change": SI

### — Supporting Information —

#### File S1

The file is available here.

#### Table S1

Description of the used predictors.

| *Code* | *Variables* |
|---|---|
| dist_pert_Fire010 | Distance to the Nearest Fire with age in $[0, 10]$ years |
| dist_pert_Fire1020 | Distance to the Nearest Fire with age in $]10, 20]$ years |
| dist_pert_Fire2050 | Distance to the Nearest Fire with age in $]20, 50]$ years |
| Age | Stand Age |
| freqCD | Frequency of Dense Conifer Forests |
| freqCO | Frequency of Open Conifer Forests |
| freqMW | Frequency of Mixed-Wood Habitat |
| freqOH | Frequency of Open Habitat |
| freqDF0_10 | Frequency of Fire disturbed Forests with age in $[0, 10]$ years |
| freqDF10_20 | Frequency of Fire disturbed Forests with age in $]10, 20]$ years |
| freqDF20_50 | Frequency of Fire disturbed Forests with age in $]20, 50]$ years |
| freqDC0_10 | Frequency of Harvest disturbed Forests with age in $[0, 10]$ years |
| freqDC10_20 | Frequency of Harvest disturbed Forests with age in $]10, 20]$ years |
| freqDC20_50 | Frequency of Harvest disturbed Forests with age in $]20, 50]$ years |

#### Table S2

The potential 3 full models.

| Models (full) | Fixed-effect without intercept |
|---|---|
| Model0 | $linear\_terms$ |
| Model1 | $linear\_terms + Age\_in^2$ |
| Model2 | $linear\_terms + Dist\_Fire\_in^2 + Age\_in^2$ |

### Figure S1

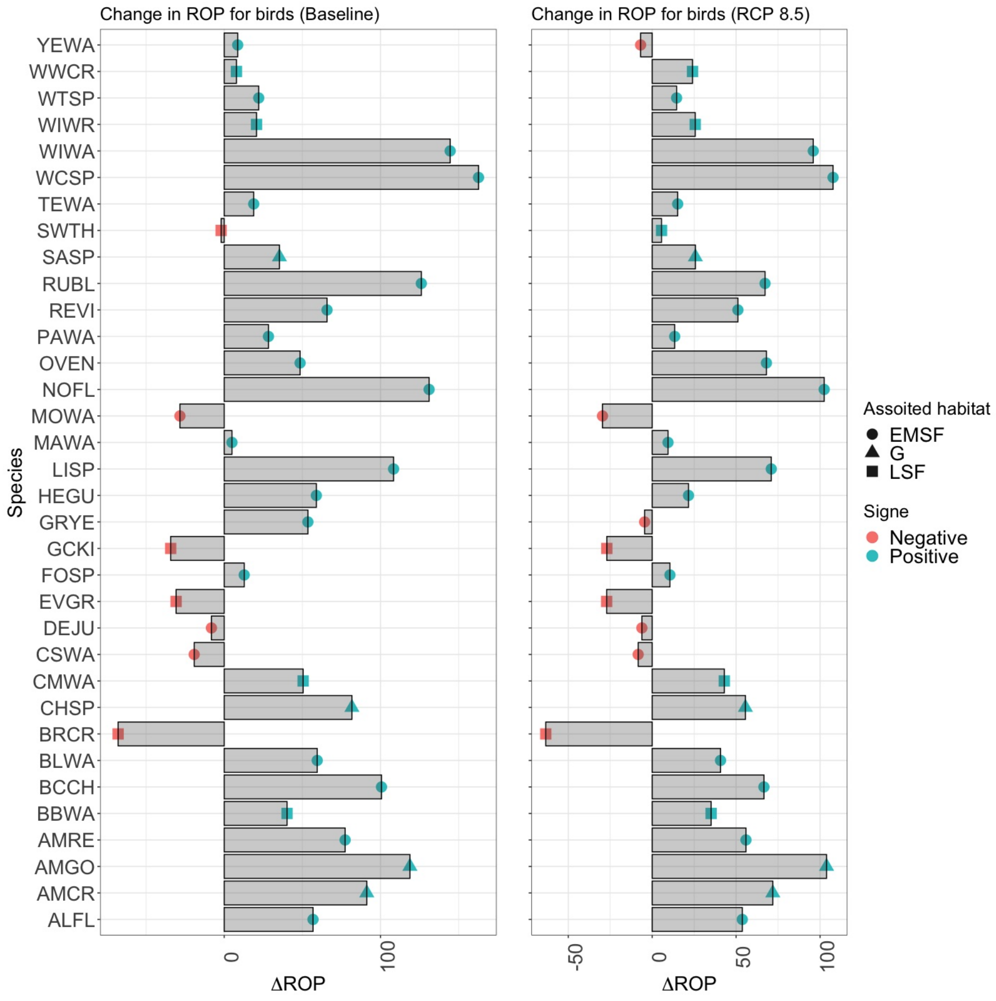


Figure S1: **Change in ROP for bird species**. Percentage of change in the regional occupancy probability under Baseline (between Baseline-NoHarvest and Baseline-Harvest) and RCP 8.5 (between RCP 8.5-NoHarvest and RCP 8.5-Harvest) scenarios for bird species. **Abbreviations**: late succession forest (*LSF*), early-to-mid succession forest (*EMSF*), Generalist (G).

### Figure S2

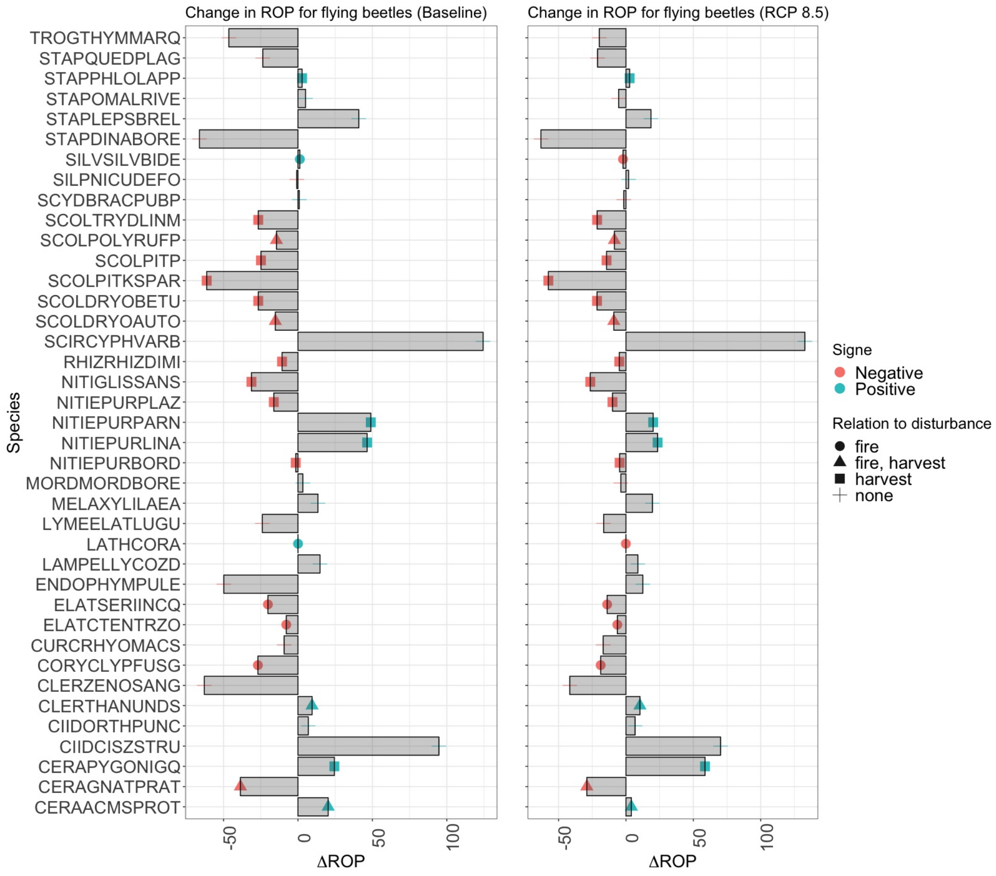


Figure S2: **Change in ROP for flying beetles species**. Percentage of change in the regional occupancy probability under Baseline (between Baseline-NoHarvest and Baseline-Harvest) and RCP 8.5 (between RCP 8.5-NoHarvest and RCP 8.5-Harvest) scenarios for flying beetles species with relation to disturbance.

### Figure S3

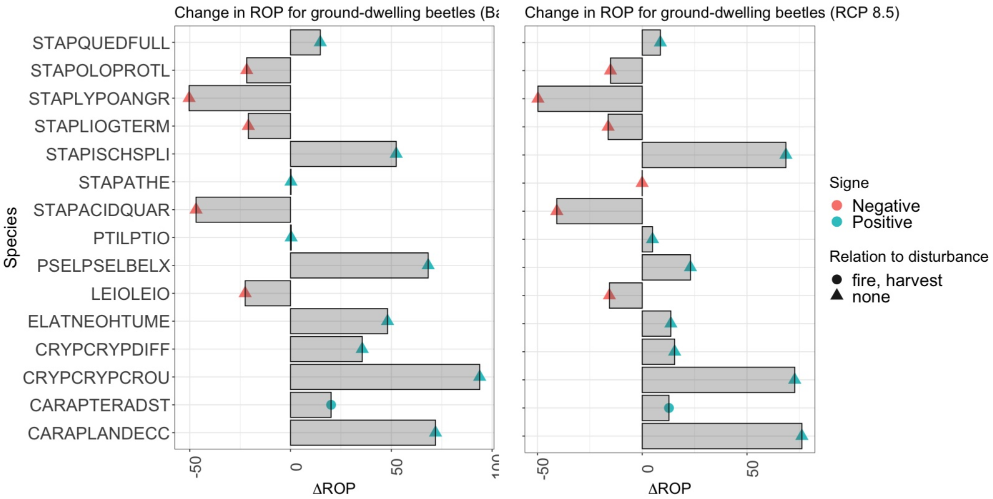


Figure S3: **Change in ROP for ground beetles species**. Percentage of change in the regional occupancy probability under Baseline (between Baseline-NoHarvest and Baseline-Harvest) and RCP 8.5 (between RCP 8.5-NoHarvest and RCP 8.5-Harvest) scenarios for ground beetles species with relation to disturbance.

### Figure S4

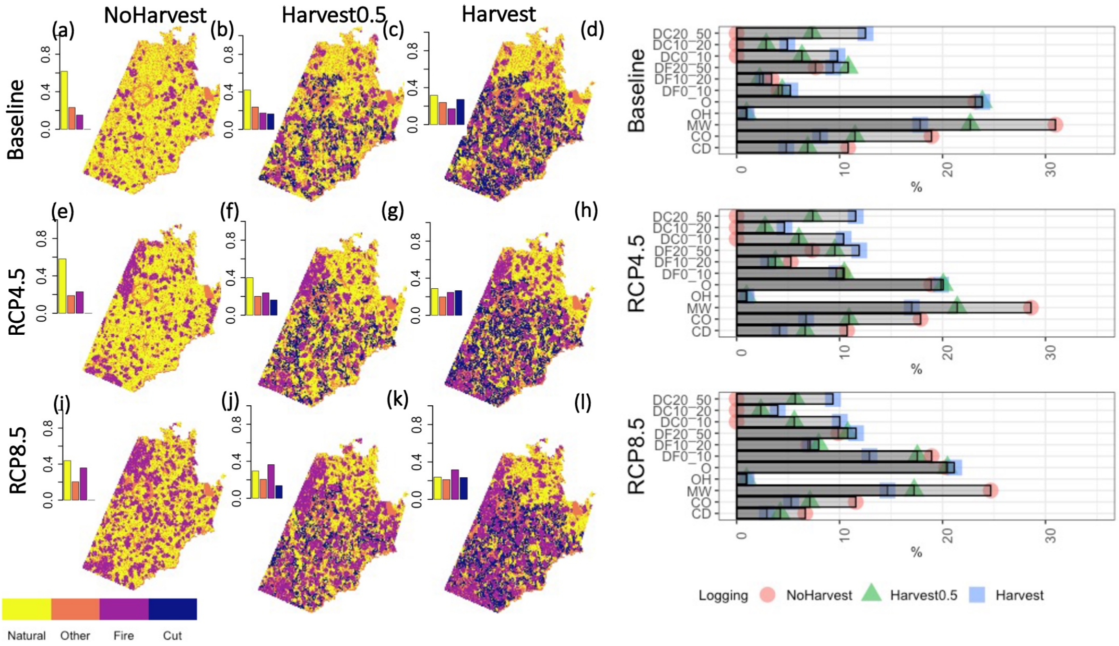


Figure S4: The distribution of the land cover under the simulated management scenarios in 2100.

**Temporary page!**

LaTeX was unable to guess the total number of pages correctly. As there was some unprocessed data that should have been added to the final page this extra page has been added to receive it.

If you rerun the document (without altering it) this surplus page will go away, because LaTeX now knows how many pages to expect for this document.
